## Supplemental Information for "Global and gene-specific translational regulation in *Escherichia coli* across different conditions"

### Supplementary Information

Includes the following items:

**1. Theoretical analysis for the distribution of correlation coefficients between mRNA level and randomly scrambled RTE.**

**2. Quantify the contribution of codon usage to two different definitions of RTE variability.**

**Supplementary Figure 1.**
Scatter plots of mRNA and ribosome levels on a per-gene basis across multiple replicates.

**Supplementary Figure 2.**
Data reproducibility of RNA-seq and Ribo-seq across multiple replicates.

**Supplementary Figure 3.**
Translational control of genes from the same polycistronic transcript.

**Supplementary Figure 4.**
QQ plot comparing the actual distribution to randomly distributed RTEs (Fig. 1B).

**Supplementary Figure 5.** Transcription and translation couple together to respond to nutrient limitations at the growth rate of 0.6 $h^{-1}$.

**Supplementary Figure 6.** No significant differences in translational regulation between the different growth rates under the same nutrient limitation.

**Supplementary Figure 7.** Global view of the 82 pathways in *E. coli* on the biplot of mean RTE and RTE variance.

**Supplementary Figure 8.** Codon ranks from random forest model are consistent with the anti-correlated codons in Fig. 5A.

**Supplementary Table 1.** Classification results of the random forest model with additional features.

**Supplementary Figure 9.** The contribution of codon usage to the classification result is robust for different definitions of RTE variability.

**Supplementary Figure 10.** Illustration of selection of two clusters of genes when comparing nutrient limiting conditions in pairs.

**Supplementary Figure 11.** Correlation between mean mRNA level and mean RTE, across different genes. Each subfigure was derived from one of the 12 conditions.

**1. Theoretical analysis for the distribution of correlation coefficients between mRNA level and randomly scrambled RTE.**

We can theoretically infer the distribution of the Spearman's rank correlation coefficients between the mRNAs and randomly scrambled RTEs of all genes. To calculate the Spearman's rank correlation coefficients, we just need to obtain the rank values of mRNA and RTE across conditions for a gene. Then we can calculate the Spearman's correlation coefficients using the popular formula

$$\rho = 1 - \frac{6\sum d_i^2}{n(n^2-1)},$$

where

$$d_i = rank(mRNA_i) - rank(RTE_i)$$

is the difference between the two ranks of mRNA and RTE for the ith condition. And n is 12 in our experiments corresponding to 12 conditions.

The RTEs were being randomly scrambled so the rank values of RTEs were randomly permutated. Since the mRNA level can be considered known and fixed in our experiments, the d is equal to a variable from a fixed sequence minus a randomly and uniformly distributed variable. Therefore, the d appears in a symmetric triangular distribution with zero mean

$$f(d) = \begin{cases} \frac{d+m}{m^2}, & -m \le x < 0 \\ \frac{m-d}{m^2}, & 0 \le x \le m \end{cases},$$

where m is the number of genes, that is 2914 in our analysis. We approximate the triangular distribution to a normal distribution, then the distribution of the sum of d squared can be considered as a $\chi^2$-distribution. Because the number of degrees of freedom (n) is large enough, it can be approximated as a normal distribution with zero mean. It should be noted that the Spearman's rank correlation coefficient is limited to [-1, 1]. Thus, the distribution of ρ is symmetric with zero mean.

**2. Quantify the contribution of codon usage to two different definitions of RTE variability.**

In addition to considering RTE variance, we used two additional evaluation indices – the Fano factor and the coefficient of variation (CV) —to test whether our conclusions are robust with respect to different definitions of RTE variability. The same analyses were performed for these two indices, and the major conclusions are consistent with those found using RTE variance. The mean codon frequencies of the top 200 and bottom 200 genes are negatively correlated (Sup Fig. 6, A-B). The addition of the feature codon frequency significantly improves the classification accuracy (Sup Fig. 6, C-D). Other features, such as translation pause motifs, do not contribute to classification accuracy (Sup Fig. 6, E-F). The contributions of mRNA and RTE to classification accuracy differ when using different definitions of RTE variability. This is because these indices have different degrees of mathematical dependence on mRNA and RTE. However, these differences do not change the main conclusion that codon usage contributes to RTE variability.

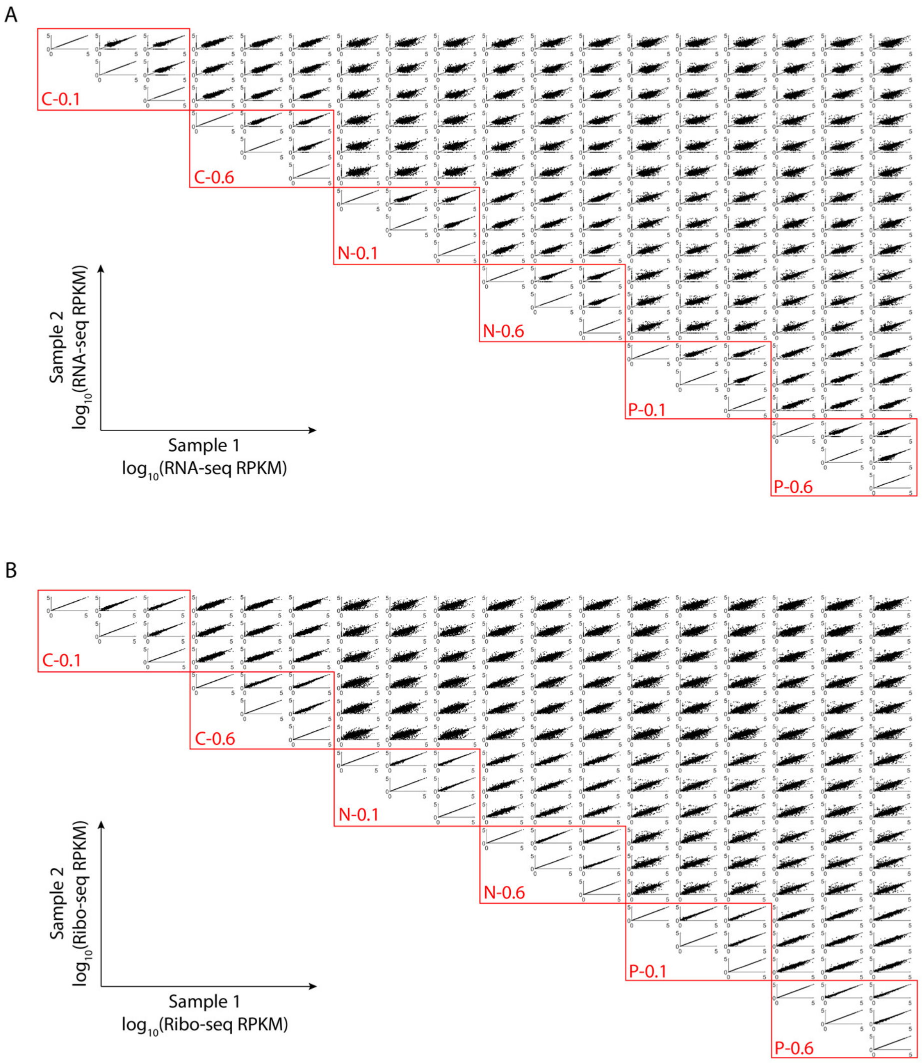


**Supplementary Figure 1.** Scatter plots of mRNA and ribosome levels on a per-gene basis across multiple replicates.

(A) Scatter plots of mRNA level per gene from RNA-seq across different replicates. Each dot represents an individual gene. In the red boxes are pairwise comparisons among three biological replicates from the same condition.

(B) Same as (A), but showing scatter plots from ribosome profiling.

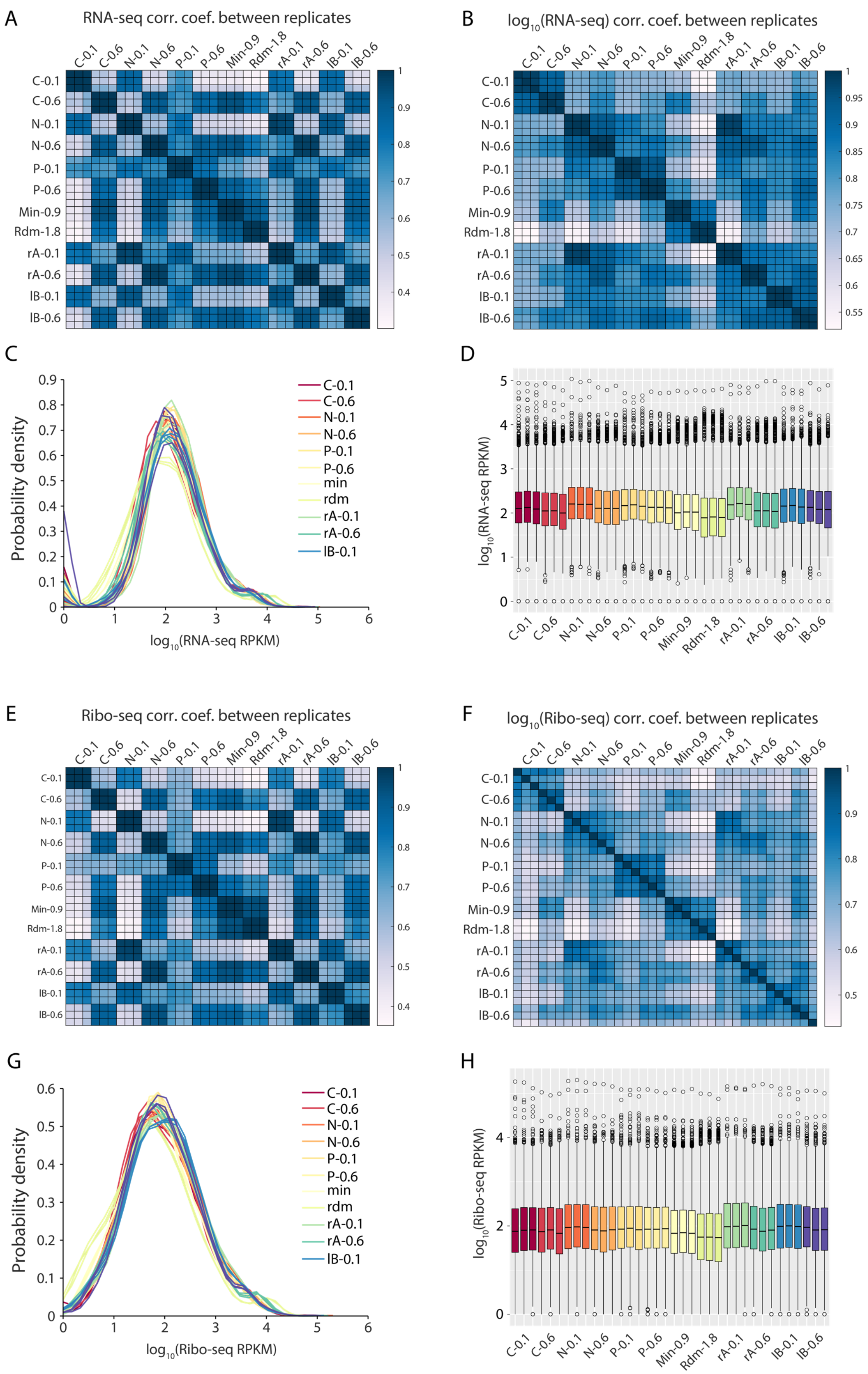
A
RNA-seq corr. coef. between replicates
C-0.1
C-0.6
N-0.1
N-0.6
P-0.1
P-0.6
Min-0.9
Rdm-1.8
rA-0.1
rA-0.6
IB-0.1
IB-0.6
B
log10(RNA-seq) corr. coef. between replicates
C
Probability density
log10(RNA-seq RPKM)
C-0.1
C-0.6
N-0.1
N-0.6
P-0.1
P-0.6
min
rdm
rA-0.1
rA-0.6
IB-0.1
D
log10(RNA-seq RPKM)
E
Ribo-seq corr. coef. between replicates
F
log10(Ribo-seq) corr. coef. between replicates
G
Probability density
log10(Ribo-seq RPKM)
H
log10(Ribo-seq RPKM)

**Supplementary Figure 2.** Data reproducibility of RNA-seq and Ribo-seq across multiple replicates.
(A) Heatmap of Pearson correlation coefficients between replicates based on RNA-seq RPKM.
(B) Heatmap of Pearson correlation coefficients between replicates based on $\log_{10}$(RNA-seq RPKM).
(C) Distribution of gene $\log_{10}$(RNA-seq RPKM) across 36 replicates. The three biological replicates for each condition are shown in the same color.
(D) Box plots of gene $\log_{10}$(RNA-Seq RPKM) across 36 replicates.
(E-H) Same as (A-D), but showing results from ribosome profiling.

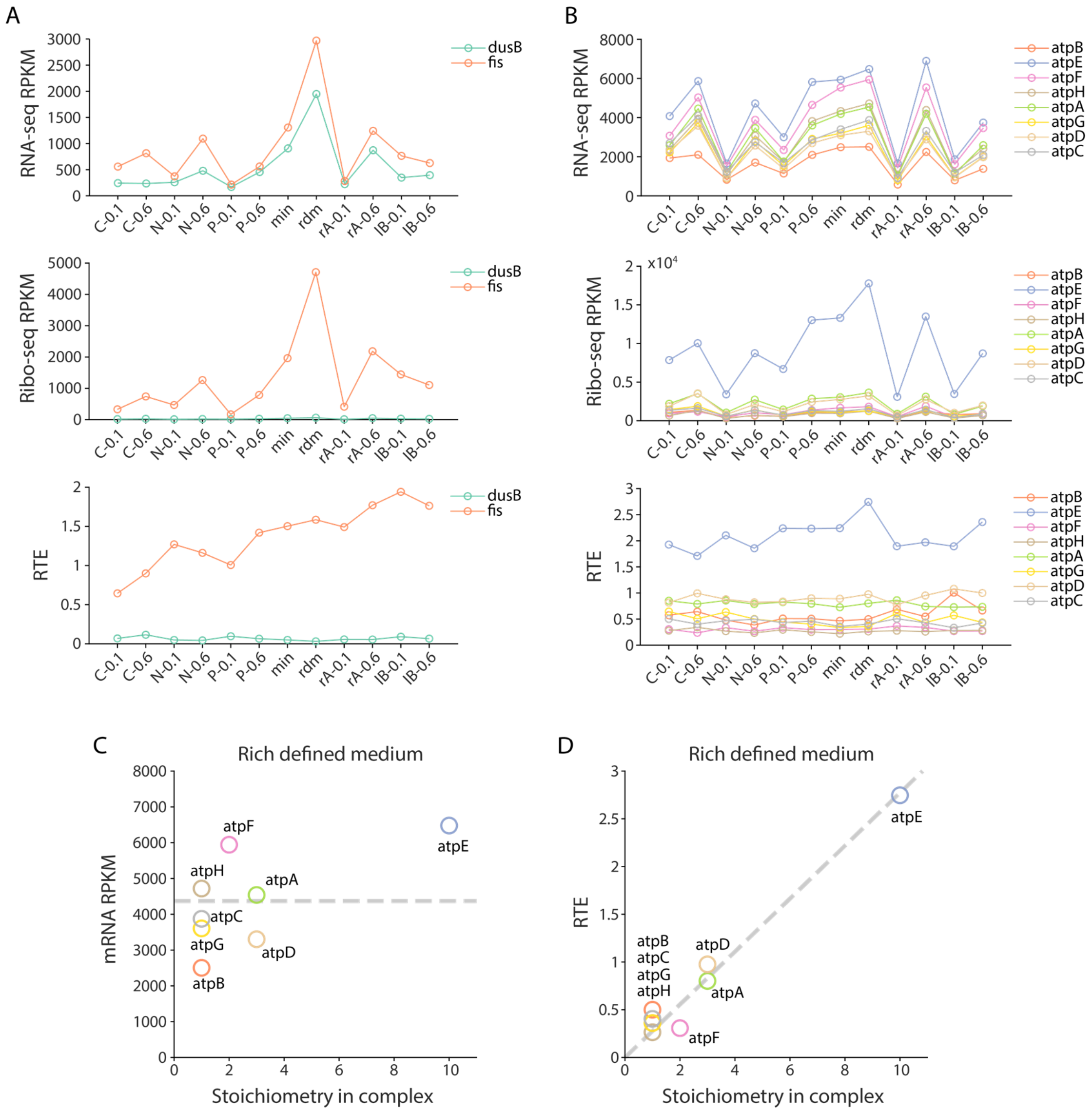


**Supplementary Figure 3.** Translational control of genes from the same polycistronic transcript. (A) Gene expression pattern across 12 conditions of two genes from the *dusB-fis* operon. The upper panel shows the RNA-seq RPKM. The median panel shows the Ribo-seq RPKM. The lower panel shows the relative translation efficiency.

(B) Gene expression pattern across 12 conditions of eight subunits from the $F_0F_1$ ATP synthase complex that are expressed from a single polycistronic transcript.

(C) mRNA level for the eight genes from the $F_0F_1$ ATP synthase complex. The average mRNA level for each gene in rich defined medium is plotted against its stoichiometry in the complex. The dashed line indicates the average mRNA level.

(D) Relative translation efficiency for the eight genes from the $F_0F_1$ ATP synthase complex. The average RTE for each gene in rich defined medium is plotted against its stoichiometry in the complex. The dashed line indicates the best-fit that crosses the origin.

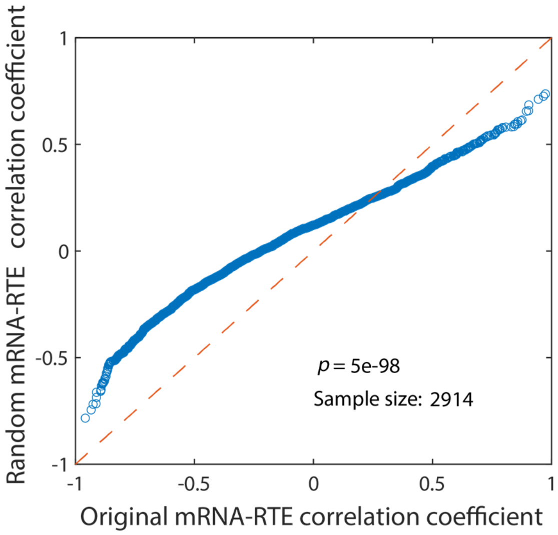


**Supplementary Figure 4.** QQ plot comparing the actual distribution to randomly distributed RTEs (Fig. 1B). Student's *t*-test was used to calculate the *p*-value.

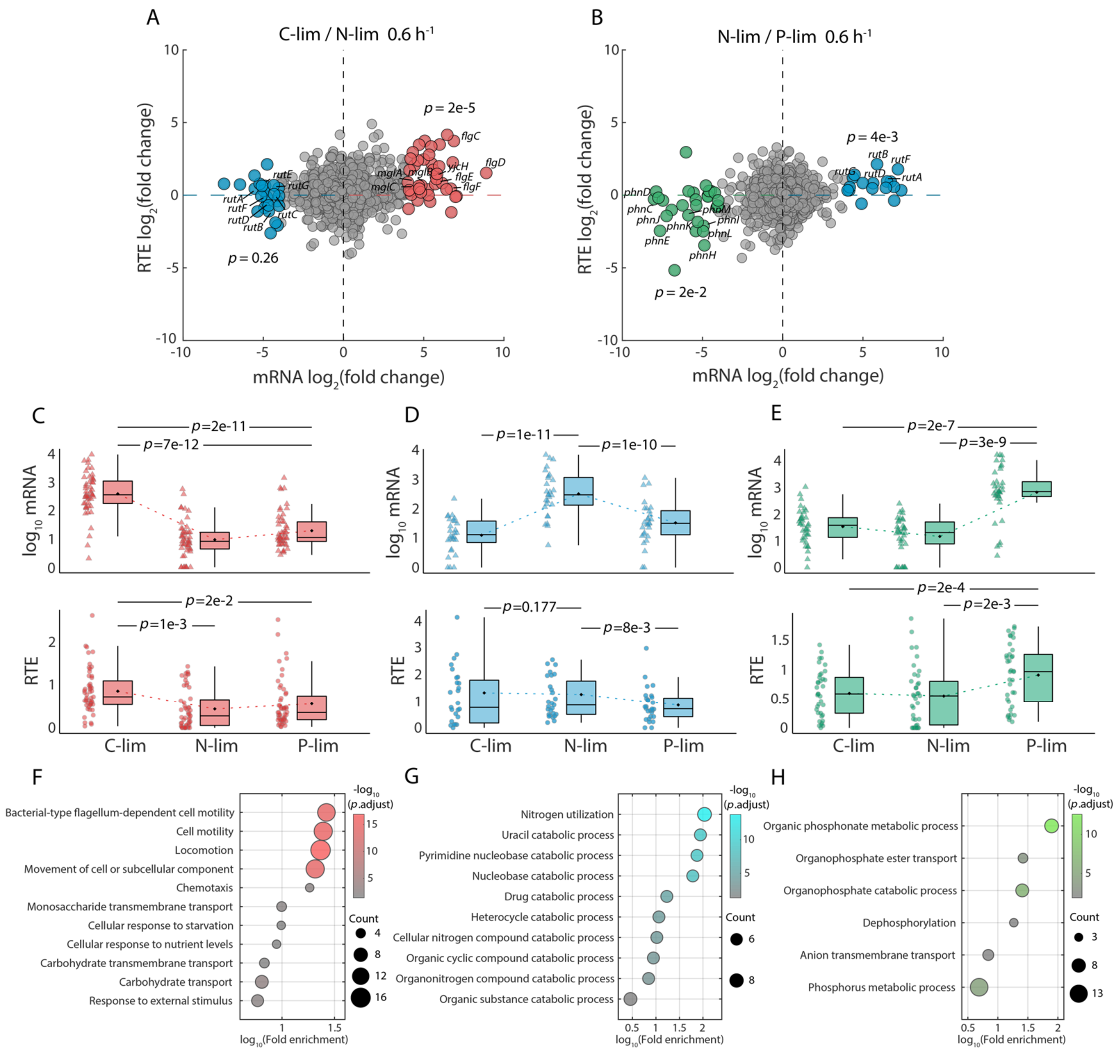


**Supplementary Figure 5.** Transcription and translation couple together to respond to nutrient limitations at the growth rate of 0.6 $h^{-1}$.

(A) Comparison of transcription changes ($log_2$ mRNA fold change, *x*-axis) and translation changes ($log_2$ RTE fold change, *y*-axis) between carbon limitation and nitrogen limitation at the growth rate of 0.6 $h^{-1}$. The averages of three biological replicates are shown. Red dots represent genes with $log_2$ mRNA fold change (C-limited / N-limited) > 4. Blue dots represent genes with $log_2$ mRNA fold change (C-limited / N-limited) < -4. *p*-value was used to test the significance of the RTE fold change between the highlighted genes and the background genes.

(B) Same as (A), but showing the change between nitrogen limitation and phosphate limitation. Green dots represent genes with $log_2$ mRNA fold change (N-limited / P-limited) < -4.

(C-E) mRNA level (upper panel) and RTE level (lower panel) of the three groups of highlighted genes in (A) and (B). The three highlighted groups of genes are upregulated under carbon (C), nitrogen (D), and phosphate (E) limitations.

(F-H) Gene ontology (GO) enrichment analysis for the three highlighted gene groups in (A) and (B). The color of the dots represents the $-log_{10}$ adjusted *p*-value, and the size represents the number of genes.

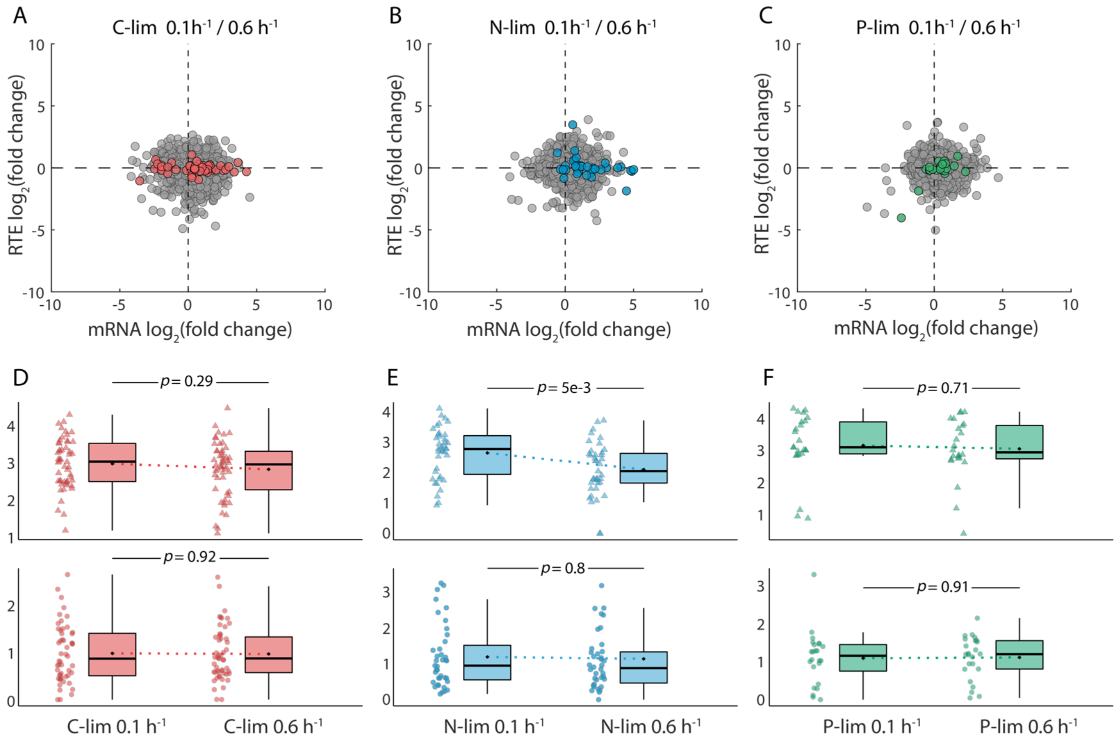


**Supplementary Figure 6.** No significant differences in translational regulation between the different growth rates under the same nutrient limitation.

(A) Comparison of transcription changes ($\log_2$ mRNA fold change, *x*-axis) and translation changes ($\log_2$ RTE fold change, *y*-axis) between the growth rate of 0.1 $h^{-1}$ and that of 0.6 $h^{-1}$ under C limitation. The averages of three biological replicates are shown. Red dots represent genes with the same color in Fig. 3A.

(B) Same as (A), but showing the comparison under N limitation and highlighting the genes in Fig. 3A with blue color.

(C) Same as (A), but showing the comparison under P limitation and highlighting the genes in Fig. 3B with green color.

(D-F) mRNA level (upper panel) and RTE level (lower panel) of the three groups of highlighted genes in (A-C). In each panel, two different growth rates are shown with the same nutrient limitation.

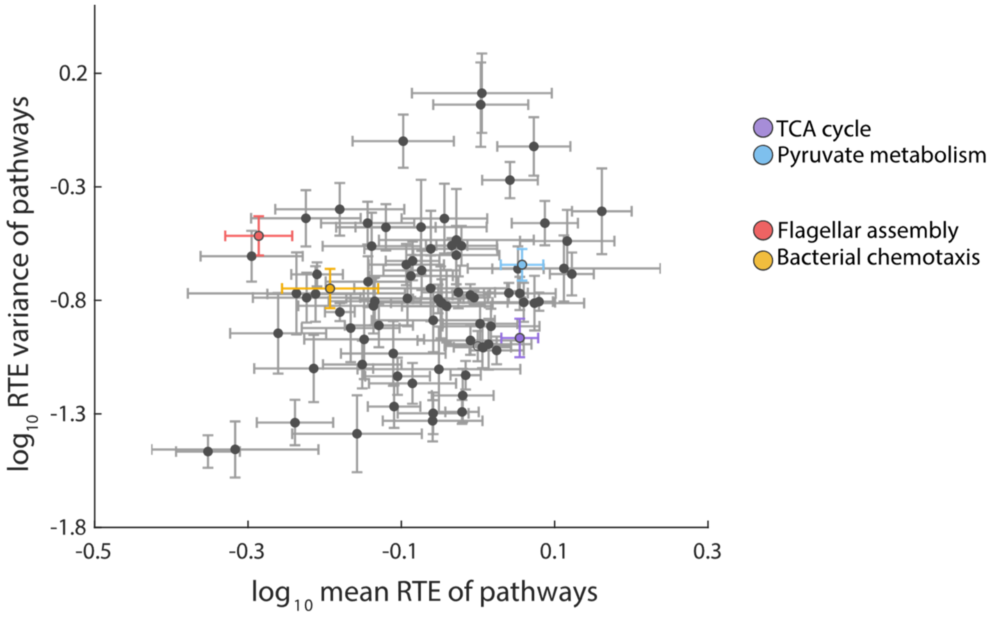


**Supplementary Figure 7.** Global view of the 82 pathways in *E. coli* on the biplot of mean RTE and RTE variance. Error bars represent s.e.m. from genes in each pathway.

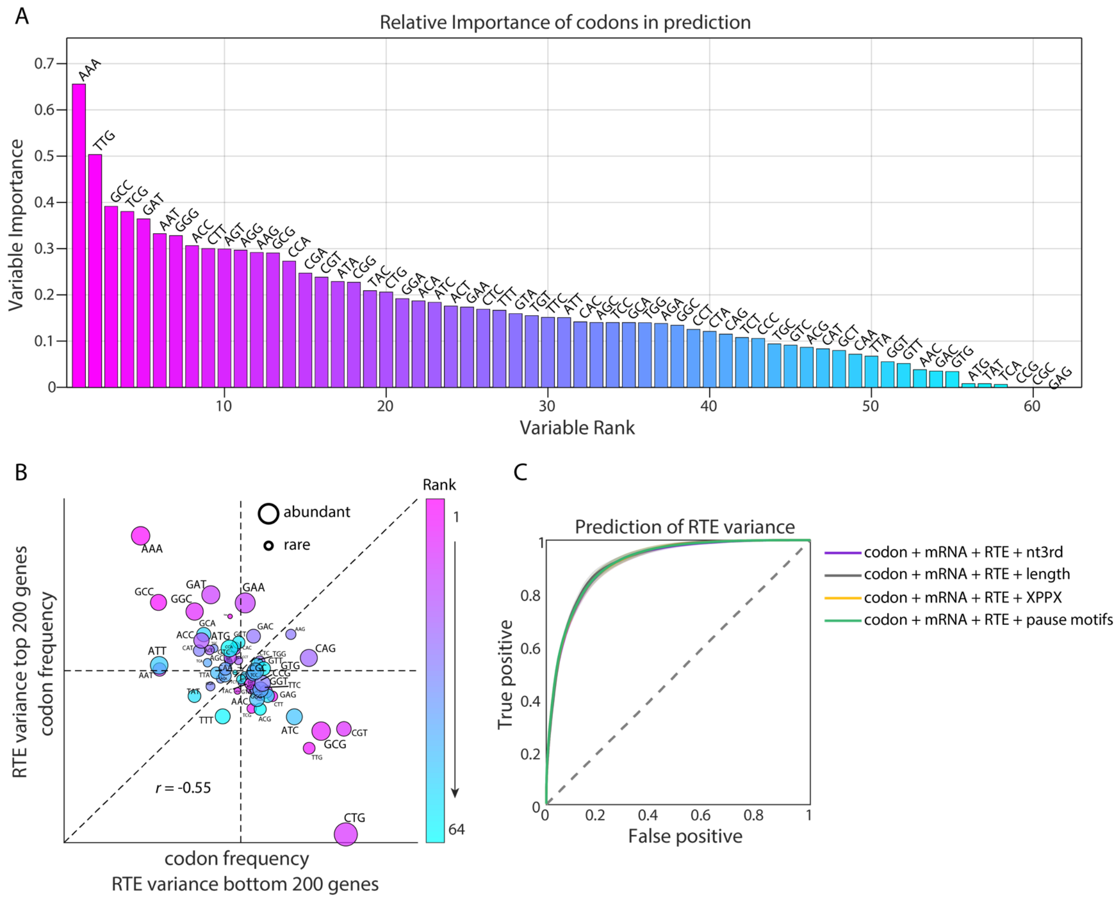


**Supplementary Figure 8.** Codon ranks from random forest model are consistent with the anti-correlated codons in Fig. 5A.

(A) The codon ranks from random forest model. The three stop codons TAA, TGA, and TAG were removed because they occur only once in each gene and do not encode amino acids.

(B) Negative correlation between the codon frequencies for the top 200 and bottom 200 genes in their RTE variances. Codons are colored by their ranks from random forest model.

(C) The ROC curves of the classification result for the variance of RTE with other features. The nt3rd stands for the distribution of the third base of the codon. The XPPX stands for XPPX motif inducing translation pause. The pause motifs stand for other codon combinations that mediate the translation pause.

| Features | Sensitivity | Specificity | Accuracy | AUC |
|---|---|---|---|---|
| All 3 + nt3rd | 84.18±2.26% | 82.82±2.23% | 83.48±1.47% | 0.91±0.01 |
| All 3 + length | 85.15±2.08% | 83.19±2.27% | 84.15±1.49% | 0.92±0.01 |
| All 3 + XPPX | 84.3±2.27% | 82.86±2.28% | 83.56±1.51% | 0.91±0.01 |
| All 3 + pause motifs | 84.4±2.11% | 82.86±2.22% | 83.62±1.43% | 0.91±0.01 |

**Supplementary Table 1.** Classification results of the random forest model with additional features.

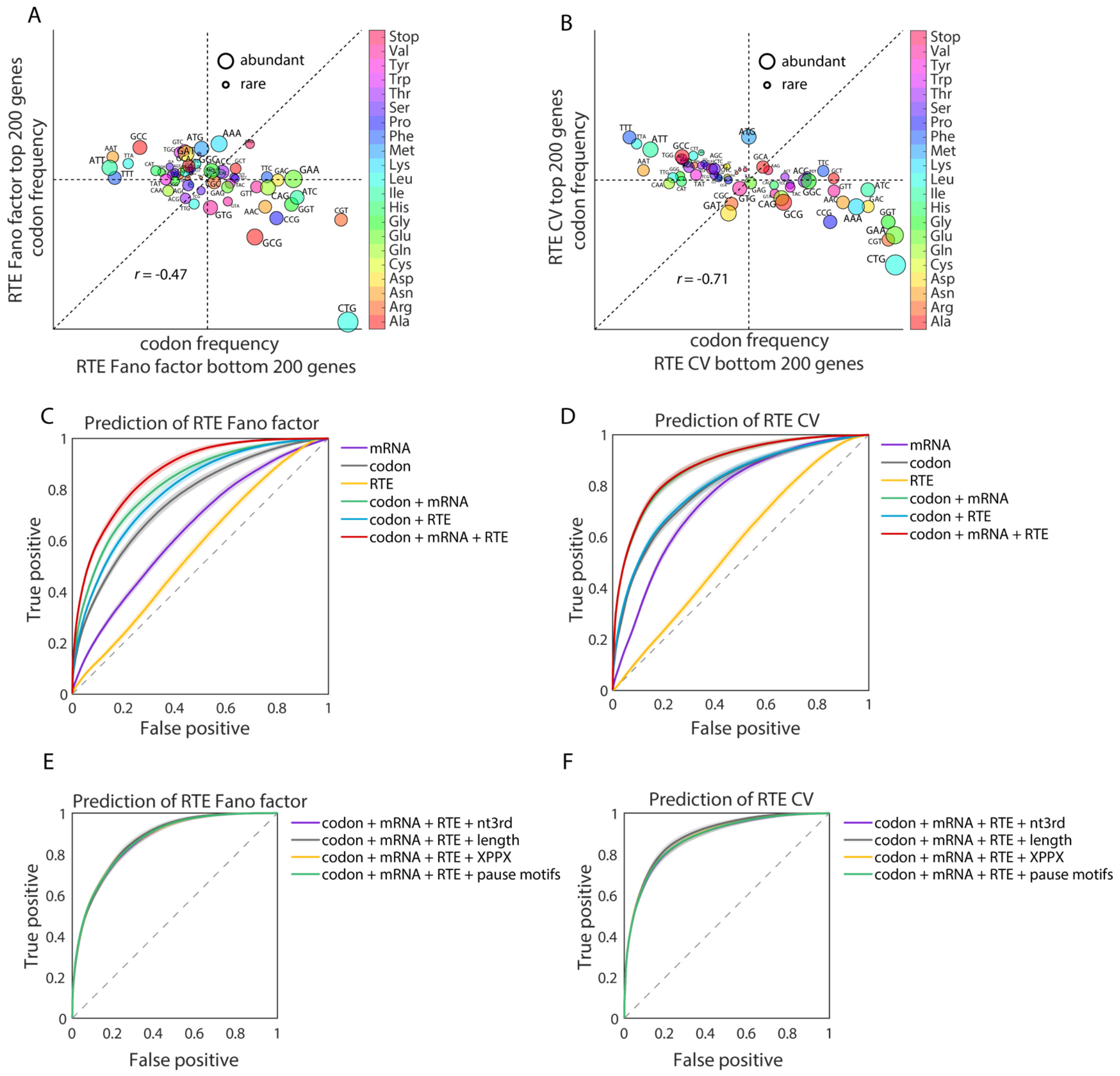


**Supplementary Figure 9.** The contribution of codon usage to the classification result is robust for different definitions of RTE variability.

(A) Correlation of the codon frequencies between the top 200 and bottom 200 genes in the Fano factor of RTE.

(B) Correlation of the codon frequencies between the top 200 and bottom 200 genes in the coefficient of variation (CV) of RTE.

(C) The ROC curves of the classification result for the Fano factor of RTE.

(D) The ROC curves of the classification result for the CV of RTE.

(E) The ROC curves of the classification result for the CV of RTE with other features.

(F) The ROC curves of the classification result for the Fano factor of RTE with other features.

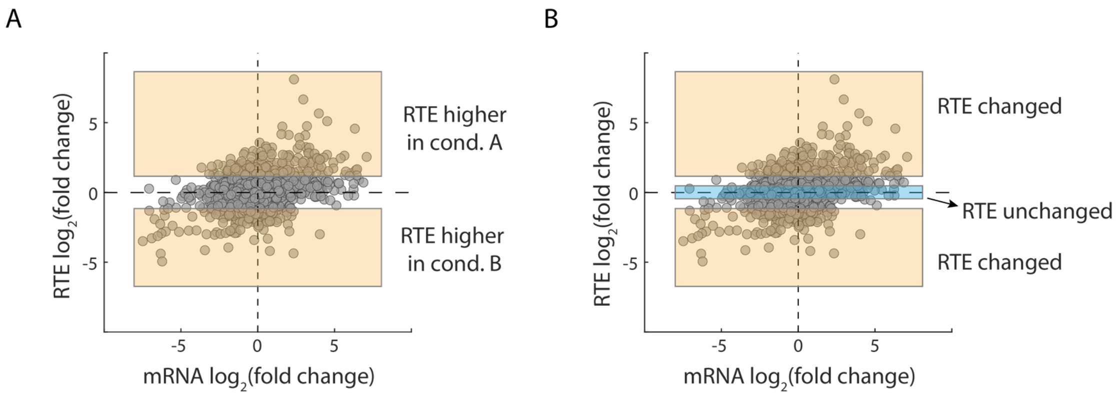


**Supplementary Figure 10.** Illustration of selection of two clusters of genes when comparing nutrient limiting conditions in pairs.

(A) When any pair of conditions A and B were compared, genes with a significantly higher RTE was filtered by $\log_2$(RTE fold change) > 1 or < -1.

(B) When any pair of conditions A and B were compared, genes with high RTE variability were filtered by $\log_2$(RTE fold change) > 1 or < -1. And the genes with low RTE variability were filtered by -0.5 < $\log_2$(RTE fold change) < 0.5.

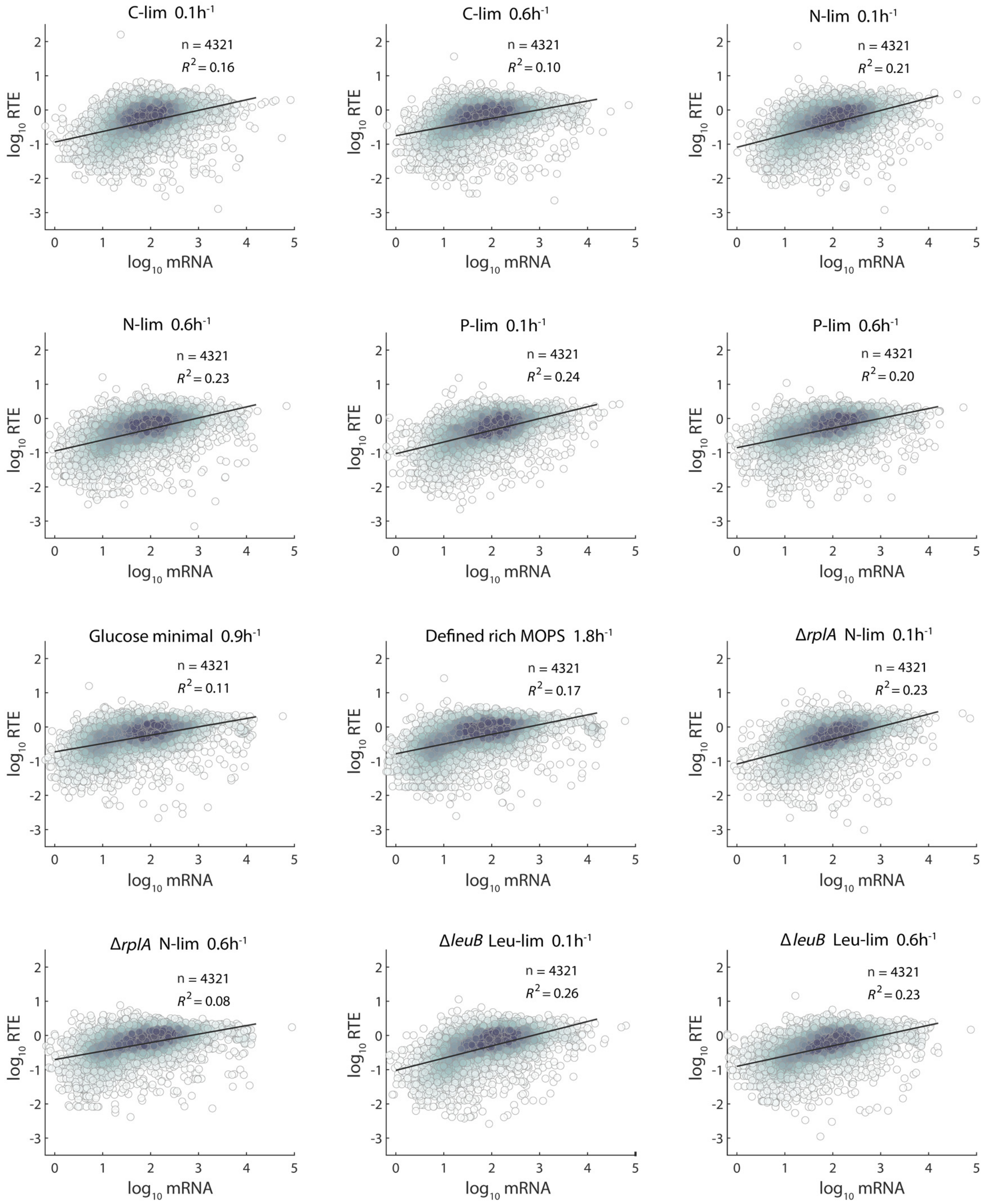


**Supplementary Figure 11.** Correlation between mean mRNA level and mean RTE, across different genes. Each subfigure was derived from one of the 12 conditions. Each dot represents one gene, and color depth depicts the density of points.
